# Storage length and temperature influence infectivity and spore yield of two common *Daphnia* parasites

**DOI:** 10.1101/354522

**Authors:** Meghan A. Duffy, Katherine K. Hunsberger

**Affiliations:** Department of Ecology & Evolutionary Biology, University of Michigan, Ann Arbor, MI 48109

**Keywords:** Metschnikowia, Pasteuria, refrigerator, freezer, burst size, spore, fitness, Daphnia dentifera

## Abstract

*Daphnia* and their parasites have emerged as a model system for understanding the ecology and evolution of infectious diseases. Two of the most commonly studied *Daphnia* parasites are the bacterium *Pasteuria ramosa* and the fungus *Metschnikowia bicuspidata*. In addition to being the focus of numerous field studies, these two parasites have been used in many laboratory experiments. However, there is little information in the scientific literature about how the conditions under which these parasites are stored influence the infectivity and yield of transmission stages (“spores”). This is problematic because such information is critical for experiment design and data interpretation.

We tested the influence of storage length (eight treatments ranging from 1 day to 1 year) and temperature (−20°C (freezer) vs. 4°C (refrigerator)) on spore infectivity and yield. We found that *Pasteuria* spores survived well at both −20°C and 4°C, and remained infective even after storage for one year. However, *Pasteuria* spore yields dropped over time, particularly at 4°C. In contrast, *Metschnikowia* spores were killed within days at −20°C. At 4°C, *Metschnikowia* infectivity declined steadily over a period of two months and, by four months, spores were no longer infective. Spore yield from *Metschnikowia*-infected hosts was not significantly impacted by storage length, but trended downwards.

Scientists working with *Pasteuria* should be aware that spore yield declines during storage, particularly in the refrigerator. Scientists working with *Metschnikowia* should be aware that it is killed by freezer storage and that, even if it is stored in the refrigerator, infectivity declines within a few months. These results might have implications for parasite distributions in the field; for example, the high sensitivity of *Metschnikowia* to freezing might help explain why it tends to be more common in deep lakes than in ponds or rock pools.

## Introduction

Studies of *Daphnia*-parasite interactions date back to at least the 1880s (Metchnikoff 1884, 1888, 1889). In the past 25 years, *Daphnia* and their parasites have emerged as a model system for understanding the ecology and evolution of infectious diseases (Ebert 2005, Lampert 2011, Cáceres et al. 2014). One strength of the *Daphnia*-parasite system is its utility in laboratory experiments, which have helped uncover factors that are relevant to the ecology and evolution of infectious diseases (Ebert 2008, Ebert 2011, Lampert 2011).

Despite being an important study system, there is surprisingly little information in the scientific literature about how lab conditions, including storage length and temperature, influence transmission stages of common *Daphnia* parasites. These transmission stages (hereafter: “spores”) are the infective stage of the parasite, enabling the parasite to move between host individuals. If lab storage conditions influence the infectivity and/or yield of parasite spores, that would affect the likelihood of an experiment succeeding. Moreover, since scientists often use infectivity and spore yield as proxies for parasite fitness, impacts of lab storage on spores could also affect estimates of parasite fitness.

Thus, understanding whether lab conditions alter spore infectivity and yield is critical for experimental design. For example, we recently carried out an experiment in which we collected spores of the common bacterial parasite, *Pasteuria ramosa*, from multiple lakes and had to store them in the lab for a year before using them in experiments. We knew that *Pasteuria* spores could remain viable in the lab for this length of time (Ebert 2005, Little et al. 2006), but had no information on whether—to maximize future infectivity and spore yield—it would be better to store them in the refrigerator (−4°C) or freezer (−20°C). For a second experiment, we wanted to kill spores of the common fungal parasite, *Metschnikowia bicuspidata*, without killing *Pasteuria* spores. We knew that *Pasteuria* spores remain viable in the freezer (Ebert 2005, Little et al. 2006), even after passing through a *Daphnia* gut (King et al. 2013). However, we did not have corresponding data on *Metschnikowia*, though, based on anecdotal observations, we suspected *Metschnikowia* spores would not remain viable in the freezer. Our two studies showed that we need more information regarding spore storage conditions to make informed decisions about appropriate experimental design.

We tested the effects of storage length and temperature on two common *Daphnia* parasites: *Pasteuria ramosa* and *Metschnikowia bicuspidata*. We stored spores for lengths ranging from 1 day to 1 year and at two temperatures (−20°C (freezer) and 4°C (refrigerator)), then assessed both their infectivity and the spore yield from infected hosts. This information will assist other scientists interested in carrying out experiments with these model systems, and might also help explain patterns of disease in natural populations.

## Methods

We tested the effects of storage length and temperature on infectivity of spores of the bacterium *Pasteuria ramosa* (“G/18” isolate) and the fungus *Metschnikowia bicuspidata* (“Std” isolate). We used infection assays in which hosts were exposed to either *Pasteuria* or *Metschnikowia*. While carrying out the experiment, one of us (KKH) noticed that the number of spores per tube seemed to decrease over time. Thus, we also did a post-hoc analysis to determine whether spore yield varied with storage length and/or temperature.

Our general procedure involved: 1) collecting *D. dentifera* that were heavily infected with either *Pasteuria* or *Metschnikowia* and putting them in microcentrifuge tubes (6-9 individuals per tube); 2) assigning each tube to a particular storage length x storage temperature treatment; 3) after the appropriate storage length, removing them from storage and grinding up the infected individuals in each tube; 4) using a hemocytometer to determine the density of spores in each tube (our measure of spore yield); 5) adding the appropriate amount of spores from a tube to a beaker containing six uninfected *D. dentifera*; 6) exposing the *D. dentifera* to the spores for 24 hours; and, finally, 7) maintaining the exposed individuals in the lab until it was possible to determine whether they were infected (our measure of infectivity).

### (a) Impact of storage length and temperature on spore infectivity

We used eight storage length treatments, ranging from 1 day to 1 year (Table 1). We used this range because sometimes we want to store spores longer term (as in the *Pasteuria* example given in the introduction), whereas at other times we want to quickly kill spores (as in the *Metschnikowia* example given in the introduction). The intermediate storage ranges were chosen because, based on prior experience with these parasites, we suspected that *Pasteuria* would live at least several months and likely a full year, whereas we suspected that *Metschnikowia* would not survive more than a few months. We used two storage temperatures (−20°C vs. 4°C), chosen to correspond with common laboratory storage conditions: in the freezer or refrigerator, respectively.

**Table 1.** Treatment combinations for the experiment. “*Pasteuria*” = *Pasteuria ramosa*; “*Metsch*” = *Metschnikowia bicuspidata*; “wks” = weeks, “mo.” = months. For some of the *Pasteuria* treatments, there were not enough spores to follow the initial plan of two low spore beakers and two high spore beakers per replicate tube. Full details can be found in the online repository for the data and code.

| Parasite | Storage temp. (°C) | Storage length | Spore doses (spores/mL) | # infected individuals per tube | # replicate tubes | # beakers per replicate tube |  | Date of exposure (month/day/year) |
| --- | --- | --- | --- | --- | --- | --- | --- | --- |
|  |  |  |  |  |  | Low spore dose | High spore dose |  |
| *Pasteuria* | 4 | 1 day | 500, 2000 | 6 | 5 | 2 | 2 | 3/14/2017 |
| *Pasteuria* | 4 | 2 wks | 500, 2000 | 6 | 5 | 2 | 2 | 3/27/2017 |
| *Pasteuria* | 4 | 4 wks | 500, 2000 | 6 | 5 | 1-3 | 2 | 4/10/2017 |
| *Pasteuria* | 4 | 2 mo. | 500, 2000 | 6 | 5 | 2 | 2 | 5/30/2017 |
| *Pasteuria* | 4 | 4 mo. | 500, 2000 | 6 | 5 | 2 | 2 | 7/11/2017 |
| *Pasteuria* | 4 | 6 mo. | 500, 2000 | 6 | 4-5 | 2-4 | 1-2 | 9/18/2017 |
| *Pasteuria* | 4 | 9 mo. | 500, 2000 | 6 | 4-5 | 2-4 | 1-2 | 12/11/2017 |
| *Pasteuria* | 4 | 1 year | 500, 2000 | 6 | 4-5 | 2 | 1-2 | 3/6/2018 |
| *Pasteuria* | -20 | 1 day | 500, 2000 | 6 | 5 | 2 | 2 | 3/14/2017 |
| *Pasteuria* | -20 | 2 wks | 500, 2000 | 6 | 5 | 2 | 2 | 3/27/2017 |
| *Pasteuria* | -20 | 4 wks | 500, 2000 | 6 | 5 | 2 | 2 | 4/10/2017 |
| *Pasteuria* | -20 | 2 mo. | 500, 2000 | 6 | 5 | 2 | 2 | 5/30/2017 |
| *Pasteuria* | -20 | 4 mo. | 500, 2000 | 6 | 5 | 2 | 2 | 7/11/2017 |
| *Pasteuria* | -20 | 6 mo. | 500, 2000 | 6 | 5 | 2 | 2 | 9/18/2017 |
| *Pasteuria* | -20 | 9 mo. | 500, 2000 | 6 | 4-5 | 2 | 2-4 | 12/11/2017 |
| *Pasteuria* | -20 | 1 year | 500, 2000 | 6 | 5 | 2 | 2 | 3/6/2018 |
| *Metsch.* | 4 | 1 day | 250, 1000 | 9 | 3 | 2 | 2 | 3/14/2017 |
| *Metsch.* | 4 | 2 wks | 250, 1000 | 9 | 3 | 2 | 2 | 3/27/2017 |
| *Metsch.* | 4 | 4 wks | 250, 1000 | 9 | 3 | 2 | 2 | 4/10/2017 |
| *Metsch.* | 4 | 2 mo. | 250, 1000 | 9 | 3 | 2 | 2 | 5/30/2017 |
| *Metsch.* | 4 | 4 mo. | 250, 1000 | 9 | 3 | 2 | 2 | 7/11/2017 |
| *Metsch.* | 4 | 6 mo. | 250, 1000 | 9 | 3 | 2 | 2 | 9/18/2017 |
| *Metsch.* | 4 | 9 mo. | 250, 1000 | 9 | 3 | 2 | 2 | 12/11/2017 |
| *Metsch.* | 4 | 1 year | 250, 1000 | 9 | 3 | 2 | 2 | 3/6/2018 |
| *Metsch.* | -20 | 1 day | 1000 | 7 | 3 | 0 | 2 | 3/14/2017 |
| *Metsch.* | -20 | 2 wks | 1000 | 7 | 3 | 0 | 2 | 3/27/2017 |

We maintain separate laboratory stock cultures of *Pasteuria* and *Metschnikowia*. We collected heavily infected *Daphnia dentifera* from these cultures on 13 March 2017; individuals were identified as heavily infected based on their increased opacity, which makes them appear bright when illuminated. We put *Pasteuria*-infected *D. dentifera* in 1.5 mL microcentrifuge tubes that were then randomly assigned to a particular temperature and storage length treatment. We then repeated the process for *Metschnikowia*-infected *D. dentifera*. The exact number of infected individuals that were added to each tube is given in Table 1.

For parasite exposures, we used two spore doses for each parasite, as we thought the lower dose would detect treatment effects at times when infectivity remained high, while the higher dose could detect differences between treatments when infectivity was low. Pilot studies indicated that *Metschnikowia* dies rapidly when stored at −20°C. Therefore, for the −20°C *Metschnikowia* treatment, we used only the high dose exposure as we expected to see no infections at the low spore dose, and included only the two shortest storage lengths, to avoid wasting spores.

We used a pestle (Fisherbrand Pellet Pestle with Cordless Motor) to grind infected *Daphnia* in one of the microcentrifuge tubes (corresponding to the treatments indicated in Table 1) and used a hemocytometer to determine the spore density in the tube. We then generated the desired spore dose by adding the appropriate volume from the tube (which contained 6-9 infected individuals, as indicated in Table 1) to a 150 mL beaker filled with 100 mL of filtered lake water (*D. dentifera* survival is greatly enhanced by culturing in filtered lake water as compared to an artificial medium). Each beaker contained six uninfected *D. dentifera* that were six days old. We used the “Mid37” genotype, which is susceptible to both the “G/18” *Pasteuria* isolate and the “Std” *Metschnikowia* isolate (Auld et al. 2014a). The number of replicate tubes is indicated in Table 1; with the exception of the *Metschnikowia* −20°C treatment (which did not have low spore beakers), a single 1.5 mL microcentrifuge tube was used to generate the spores for low spore dose and high spore dose beakers, as indicated in Table 1 in the “# beakers per replicate tube” columns.

We exposed *D. dentifera* to spores at 20°C in 16:8 light:dark (consistent with previous studies; e.g., Auld et al. 2014a). After 24 hours of spore exposure, the six *D. dentifera* individuals in each beaker were transferred to beakers containing filtered lake water that did not contain spores. Individuals were fed 10,000 cells/mL of a nutritious green alga, *Ankistrodesmus falcatus*, during parasite exposure and 20,000 cells/mL otherwise; we used a lower food dose during parasite exposure because it results in higher infection levels at a given spore dose (Hall et al. 2007). We added food to beakers four days/week; feeding four vs. seven days a week has no impact on our ability to determine whether individuals are infected, and reduces labor associated with feeding. Individuals were maintained in the lab until it was possible to determine whether they had been successfully infected (approximately 28 days for *Pasteuria* and 10-11 days for *Metschnikowia*). Infections were diagnosed by observing individuals under a dissecting microscope; as indicated earlier, infected hosts are much more opaque, with spores filling their hemolymph.

We carried out two analyses of the infectivity data, both using generalized linear models (glms) with a binomial error structure. First, we analyzed the *Pasteuria* data using a model that included dose, storage temperature, storage length (in days), and their interactions as independent variables, and a matrix of uninfected and infected hosts (per individual beaker) as the response variable. For these analyses, we used storage length as measured in days, using the exact lengths calculated from the difference between 13 March and the day of exposure given in Table 1. Second, we analyzed the data from the *Metschnikowia* 4°C treatment on its own, as we only had two storage lengths for the −20°C treatment. This model included dose, storage length (in days), and their interaction as independent variables, and a matrix of uninfected and infected hosts (per individual beaker) as the response variable. In some cases, the number of days did not precisely match to the treatment label (in months or years) due to logistical constraints (e.g., the “1 year” treatment was actually 358 days). The exact treatment lengths were as follows: 1 day (1 day), 2 weeks (14 days), 4 weeks (28 days), 2 months (78 days), 4 months (120 days), 6 months (189 days), 9 months (273 days), 1 year (358 days).

### (b) Variation in spore yield over time

We also used glms to analyze the spore yield data, which was obtained with the hemocytometer counts described above. We analyzed data on *Pasteuria* spore yield using a model that included dose, storage length (in days), storage temperature, and their interactions. For the analysis of *Metschnikowia* spore yield in the 4°C treatment, we used a model that included dose, storage length (in days), and their interaction as independent variables. For both analyses, the dependent variable was ln(spores per infected *Daphnia*).

All analyses were done in R 3.5.0. All data and code can be found on github: https://github.com/duffymeg/sporestorage.

## Results

*Pasteuria* spores remained infective even after 1 year of storage (Figure 1; Table 2). Post-hoc analyses indicate that the increase in infectivity with storage length was significant for the high spore treatment at both temperatures, and for the low spore treatment at 4°C.

**Figure 1.**
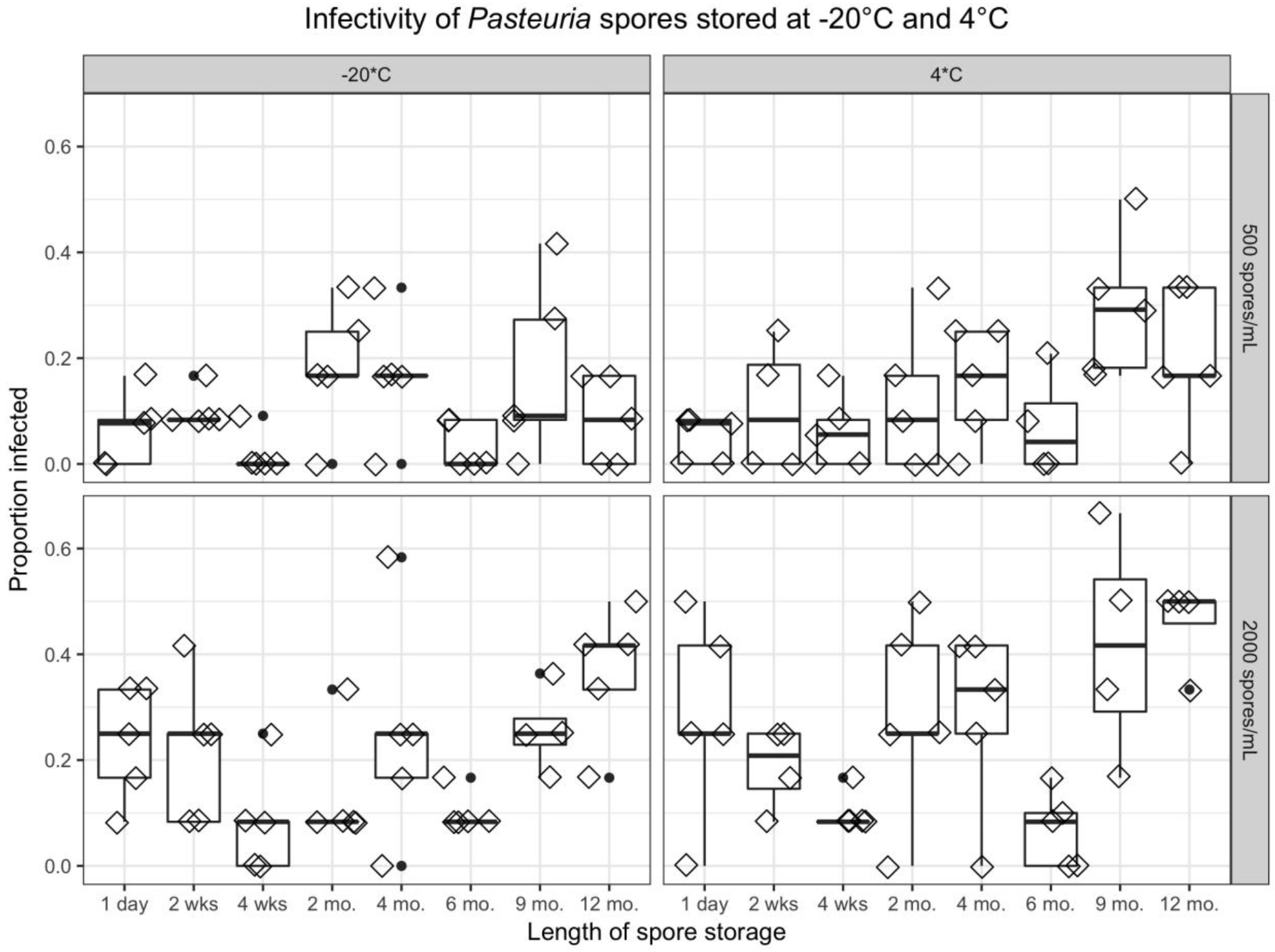
Spores of the bacterium *Pasteuria ramosa* remained infective even after a year in storage. Top row: Low spore dose treatment (500 spores/mL); bottom: high spore dose treatment (2000 spores/mL). Left: −20°C treatment (freezer); right: 4°C treatment (refrigerator). Spores remained infective in all of these treatments. Post-hoc analyses indicated that infectivity increased in the high spore treatment at both temperatures and in the low spore treatment at 4°C (*p* < 0.015 in all three treatments).

**Table 2.** Results of statistical analysis of factors influencing infectivity of *Pasteuria* spores.

| Effect | **Z** | **p** |
| --- | --- | --- |
| Storage temperature | -0.63 | 0.529 |
| Storage length | 0.83 | 0.408 |
| Dose | 2.06 | **0.040** |
| Storage temperature * storage length | 2.01 | **0.044** |
| Storage temperature * dose | 1.10 | 0.273 |
| Storage length * dose | 0.93 | 0.353 |
| Storage temperature * storage length * dose | -1.43 | 0.153 |

Spore yield from *Pasteuria*-infected hosts declined over time during the study, especially at 4°C (Figure 2; storage length: t = −2.84, *p* = 0.006; storage temperature: t = −2.04, *p* = 0.045; interaction: t = −2.50, *p* = 0.014). Even in the −20°C treatment, there was a significant drop in spore yield with increased storage length (t = −2.69, *p* = 0.011). It is possible that the increase in infectivity seen above (Figure 1) and the drop in spore yield (Figure 2) are both explained by non-viable spores degrading over time to the point where we could not visually identify them as spores.

**Figure 2.**
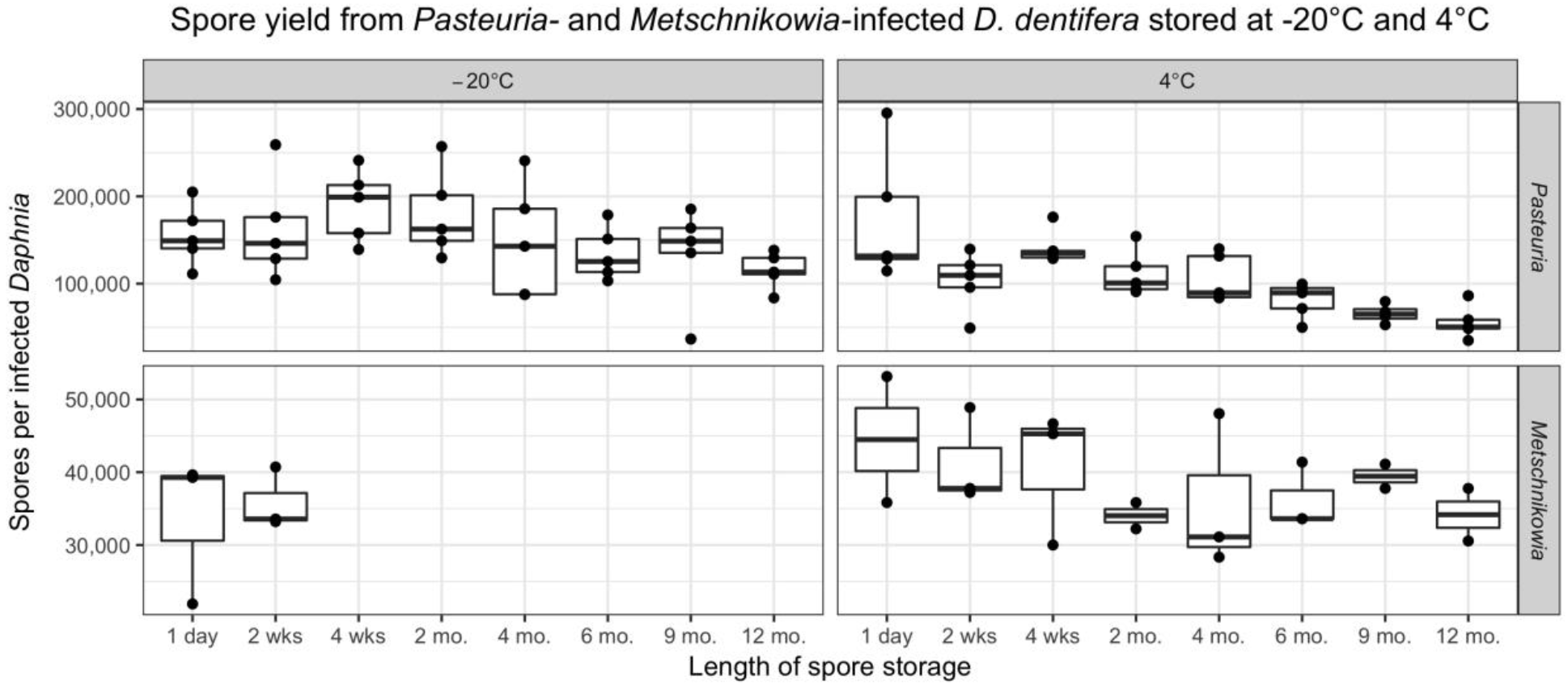
Spore yield from *Pasteuria*-infected hosts (top row) declined with increasing storage length, especially at 4°C (top right panel). Spore yield from *Metschnikowia*-infected hosts stored at 4°C (bottom right panel) did not change significantly over time, but trended downwards.

Infectivity of *Metschnikowia* spores was strongly influenced by temperature and length of storage (Figure 3). Contrary to *Pasteuria, Metschnikowia* spores died rapidly in the freezer. Only one individual became infected out of the 36 that were exposed to a high dose of *Metschnikowia* spores that had been stored at −20°C for 24 hours; in contrast, 17 of the 36 individuals exposed to spores that had been stored at 4°C for 24 hours became infected. No individuals became infected when exposed to *Metschnikowia* spores that had been stored at −20°C for two weeks. The infectivity of *Metschnikowia* spores that had been stored at 4°C declined over the first two months of storage and, by four months, spores were no longer infective. In our analysis of the *Metschnikowia* 4°C treatment data, only storage length significantly impacted infectivity (dose: z = 0.280, *p* = 0.780; storage length: z = −3.58, *p* = 0.0003; interaction: z = 0.840, *p* = 0.401).

**Figure 3.**
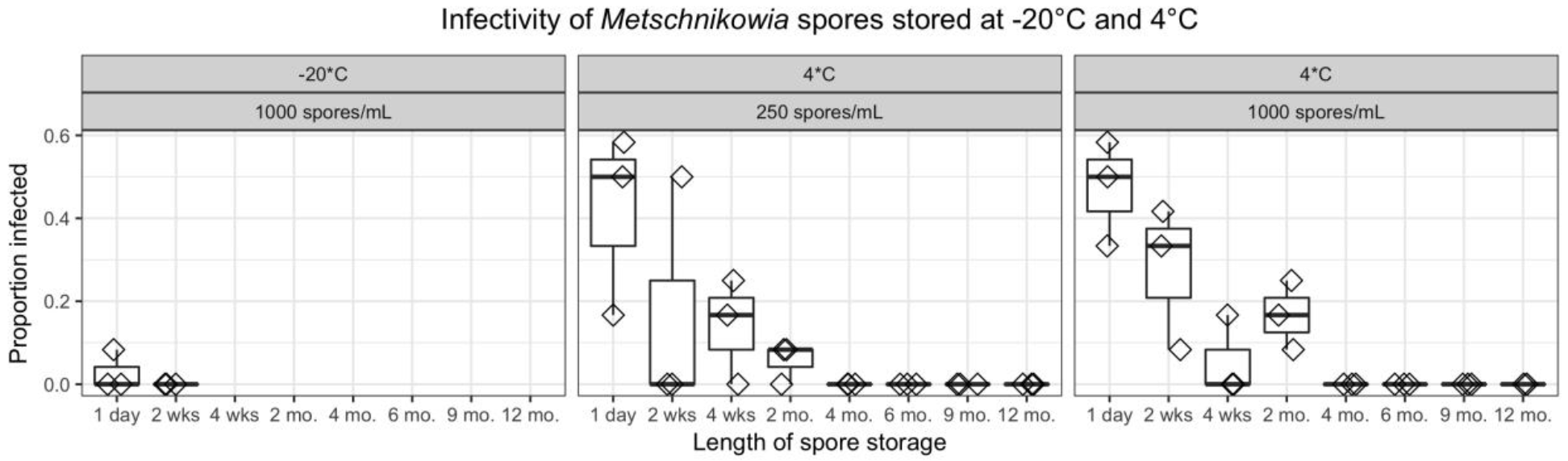
Spores of the fungus *Metschnikowia bicuspidata* rapidly lost infectivity. *Metschnikowia* spores were killed within days at −20°C (left panel) and infectivity declined sharply over 2 months at 4°C (center and right panels).

At 4°C, spore yield from *Metschnikowia*-infected hosts did not change significantly with storage length (t = −1.25, *p* = 0.229; Figure 2). However, the trend was similar to that in *Pasteuria*-infected hosts (Figure 2), and the lack of significance may simply reflect lower power based on having only three replicate tubes for *Metschnikowia* vs. five for *Pasteuria* (Table 1).

## Discussion

Infectivity and spore yield of two commonly studied *Daphnia* parasites were strongly influenced by storage length and temperature. Spores of the bacterium *Pasteuria ramosa* survived and remained infective for at least a year in both the refrigerator and the freezer. However, spore yield from *Pasteuria*-infected hosts unexpectedly declined over time, especially in the refrigerator. Spores of the fungus *Metschnikowia bicuspidata*, on the other hand, were rapidly killed by freezing; when stored in the refrigerator, *Metschnikowia* infectivity declined sharply over the first two months of storage and, by four months, spores were no longer infective. Spore yield of *Metschnikowia*-infected hosts trended downwards, but was not significantly impacted by storage length. Thus, when using these parasites in lab experiments, protocols should take into account the effects of storage length and temperature on spore infectivity and yield.

The differential response of *Pasteuria* and *Metschnikowia* spore infectivity to storage has important implications for experimental design (Figures 1 and 3). While *Pasteuria* spores can be stored in the lab for relatively long periods (months to years) before being used to infect hosts, *Metschnikowia* spores lose their infectivity over a period of weeks to months. Because of this, our lab always seeks to use *Metschnikowia* spores that are less than one month old for experiments. This presents challenges for experiments exposing field-collected hosts to field-collected spores. Such experiments require growing enough individuals of each host genotype in the lab prior to spore exposure. Ideally, these hosts would be grown for several generations under standardized conditions to control for maternal effects (i.e., influences of the mother’s environment on an offspring’s phenotype). However, experiments with field-collected hosts and spores must trade leaving more time to grow more host individuals against the rapid loss of infectivity of the field-collected spores.

The rapid death of *Metschnikowia* in the lab raises questions about how spores survive in the environment between epidemic outbreaks. *Metschnikowia* dynamics in lake populations are strongly seasonal, with disease outbreaks occurring in late summer and autumn (Duffy et al. 2009). Moreover, in the *Daphnia-Metschnikowia* system that has been best studied, the host, *Daphnia dentifera*, is dormant in the sediment for several months of the year. Thus, *Metschnikowia* is presumably able to remain infective in sediments longer than in microcentrifuge tubes in the lab. Indeed, it is possible to take sediment from a lake that had an epidemic in a previous year and use it to start epidemics in whole-water column enclosures in a lake that did not have an epidemic (A.J. Tessier, personal communication).

*Metschnikowia*’s intolerance of freezing might help explain patterns of disease in natural populations. While *Metschnikowia* has been found in ponds (Green 1974, Codreanu & Codreanu-Balcescu 1981), its prevalence in those habitats tends to be low (<4%) (Stirnadel & Ebert 1997, Goren & Ben-Ami 2013). In stratified lakes, on the other hand, *Metschnikowia* prevalence can be quite high, reaching 20% – 40% (Hall et al. 2011, Strauss et al. 2016). *Metschnikowia*’s intolerance of freezing might be a factor driving this pattern, as the sediments in ponds are much more likely to freeze. However, it is likely that other factors—such as different predation environments and stratification itself—also influence the relative prevalence in lakes vs. ponds (Auld et al. 2014a).

Spore yield from infected hosts is an important component of parasite fitness and, as a result, is frequently measured in studies of these two parasites (e.g., Jensen et al. 2006, Duffy et al. 2011, Auld et al. 2014a, Auld et al. 2014b). Our results suggest that the spore yield from infected hosts decreases over time, especially if the hosts are stored in the refrigerator (Figure 2). Ideally, spore counts from infected hosts should be done as soon after collection as possible. However, if spore counts cannot be done quickly (within days to weeks), our results suggest that samples should be stored in the freezer until they can be counted to reduce spore loss (for *Metschnikowia*, this should only be done if the spores are not needed for future infections). Even with freezer storage, spore yield will decline over time, so it is especially important to intersperse individuals from different treatments when counting samples, rather than counting all individuals from a single treatment before moving on to the next.

Our experiment used one isolate of *Pasteuria* and one isolate of *Metschnikowia*. It is possible that genotypes differ in their ability to tolerate storage under lab (and field) conditions. We hypothesize that such variation is more likely in *Pasteuria* than in *Metschnikowia*, since *Pasteuria* shows substantial variation (Carius et al. 2001, Mouton & Ebert 2008, Auld et al. 2012), whereas *Metschnikowia* shows strikingly little phenotypic and genotypic variation (Duffy & Sivars-Becker 2007, Wolinska et al. 2009, Searle et al. 2015). However, whether either parasite contains genetic variation related to tolerance of storage conditions remains to be tested.

*Daphnia* and their microparasites have emerged as an important study system for understanding the ecology and evolution of infectious diseases (Ebert 2005, Lampert 2011, Cáceres et al. 2014). Our study provides valuable information on how storage length and temperature influence spore infectivity and yield from infected hosts. When designing experiments, scientists should take into account that spore yield of *Pasteuria* declines over time, especially at 4°C, and that *Metschnikowia* spores are killed rapidly at −20°C and die within weeks to months at 4°C.

## Acknowledgments

This work was supported by the US National Science Foundation (DEB-1353806). We thank Carla Cáceres for helpful discussions and Sarah Boon for comments on this manuscript.

